# Using a lanthanide tagged nanobody to visualise Glial fibrillary acidic protein by secondary ion mass spectrometry

**DOI:** 10.64898/2026.10.22.758419

**Authors:** Clara Dieterle, Siva Ramadurai, Matija Lagator, Lauren Eyssen, Raymond J. Owens, Nicholas P. Lockyer, Hua Tian, Felicia Green

**Affiliations:** Rosalind Franklin Institute, Building R113 Rutherford Appleton Laboratory, Harwell Campus, Didcot, OX11 0QX, UK; Department of Chemistry, Photon Science Institute, The University of Manchester, Oxford Road, Manchester, M13 9PL, UK; Structural Biology Division, Centre for Human Genetics, University of Oxford, Oxford, UK; Redox Center, Department of Pediatrics, Columbia University Irving Medical Center, New York, USA

**Keywords:** nanobodies, mass spectrometry imaging, lanthanide labelling, multiplex imaging, tissue imaging, glial fibrillary acidic protein, immunofluorescence microscopy, secondary ion mass spectrometry

## Abstract

Lanthanide-labelled antibodies have been widely used for multiplex imaging of proteins directly from tissue using two approaches, imaging mass cytometry (IMC) and variants of secondary ion mass spectrometry (SIMS). Here we show the same approach can be taken using SIMS, but utilising lanthanide-labelled nanobodies instead. Nanobodies offer several advantages over conventional antibodies for antigen tagging, owing to their small size (12-15 kDa), high stability, and solubility. Their reduced molecular size enables greater diffusion through tissue, facilitating access to target sites and allowing them to reach regions that may be less accessible to larger antibodies. This enhanced accessibility can promote more effective localisation and increase binding to target antigens. In the work presented here, we produced a lanthanide-tagged nanobody and assessed its efficacy and compatibility with mass spectrometry imaging (MSI). We showed successful detection of Glial fibrillary acidic protein (GFAP) in tissue by correlating MSI of a lanthanide-tagged anti-GFAP nanobody with immunofluorescence microscopy imaging (IF). The use of nanobodies is an excellent alternative to current antibody tagging methods for SIMS and IMC.

## Introduction

Lanthanides are not endogenously found in human tissue which makes them excellent reporters with naturally low background signal in tissue when analysed using MSI techniques [1]. By conjugating lanthanides to antibodies, proteins of interest can be immunolabeled and analysed by MSI with low background noise and high sensitivity [2].

Lanthanide-labelled antibodies were first used to image proteins of interest within a biological tissue of Alzheimer’s disease with Laser Ablation-Inductively Coupled Plasma-Mass Spectrometry (LA-ICP-MS) [3]. Since then, a specifically designed instrument, Imaging Mass Cytometry (Standard Biotools™), has been developed and applied to image proteins in tissue across several scientific fields as described in the review by Thai *et al*.[2]. This method has been shown to be capable of multiplex spatial mapping of proteins with up to 40 lanthanide-coupled antibodies imaged [4] and with spatial resolutions of ∼ 2 µm and an acquisition rate of 250 pixels per second [2].

An alternative method to image the lanthanide-labelled antibodies is SIMS, with a specific instrument developed solely for this purpose called Multiplexed Ion Beam Imaging (MIBI) [2], [5]. This instrument uses an oxygen duoplasmatron primary ion beam, providing a system capable of identifying up to 100 different metal-isotopes simultaneously with a spatial resolution of ∼ 0.4 µm [6]. According to Lee *et al*. [7], MIBI is more sensitive than IMC and images at a higher spatial resolution with a faster acquisition rate, however it is no longer commercially available.

Apart from these specialised instrumentations (IMC and MIBI), the lanthanide-multiplexing approach has been adopted for ToF-SIMS usage which allows multi-omics imaging using different primary ion beams on the same tissue sample [8]. A Gas Cluster Ion Beam (GCIB) or other polyatomic beams such as C_60_ can be used initially to image lipids and metabolites. This can be followed by staining with lanthanide-conjugated antibodies for protein imaging. These imaging modalities can then be colocalised for multi-omics analysis [8], [9]. Furthermore, Electron Cyclotron Resonance (ECR) ion sources have been used in research since the late 1980s, where they were commonly applied for semiconductor etching and deposition [10]. ECR ion sources are now also used for SIMS analysis by delivering an intense beam of singly charged ions. These ion beams are generated by the ionisation of pure noble gases, such as monoatomic xenon (X^+^) and argon (Ar^+^). The advantage of ECR primary ion beams for SIMS analysis is their ability to achieve high spatial resolution (down to 300 nm) while being operated at high beam currents [11]. While higher beam currents enable faster image acquisition, they also lead to greater fragmentation of organic molecules. Consequently, ECR primary ion beams are well suited for imaging inorganic ions such as lanthanides and low-molecular weight species but are less suitable for the analysis of intact high-mass organic molecules.

Previously, lanthanide tags have only been conjugated to antibodies, RNA-probes or nanoparticles [12], [13]. In this paper, we use the same approach to conjugate in-house generated nanobodies to lanthanide tags. Nanobodies have higher water solubility, thermostability, and smaller size (∼15 kDa) in comparison to antibodies (∼150 kDa) [14]. For these reasons nanobodies are hypothesised to diffuse more readily in tissue and bind more efficiently to target proteins [14], [15], [16], [17], [18]. This would, in turn, result in increased signal intensity and spatial resolution for ToF-SIMS analysis of the associated lanthanides and would thus provide a more reliable and accurate mapping of the tagged chemical species.

## Methods

### Preparation of mouse brain sections

Fresh frozen wild type (WT) mouse brain was purchased from the Mary Lyon Centre at MRC Harwell and stored at -80 °C until cryo-sectioning. The tissue was cut between -16 °C and - 20 °C into 5 or 10 μm thick sections using the Leica CM1860 UV cryostat. Thicker sections (10 µm) were used initially for IF imaging as well as SIMS analysis using a C_60_ primary ion beam (Figure 1, 2 and 5). For SIMS analysis with the ECR primary ion beam 5 µm sections were used to reduce charging effects (Figure 4). For IF imaging, sections were thaw-mounted onto Superfrost™ plus adhesion microscope slides (Epredia) using heat from a thumb swiftly on the back of the slide until the tissue adhered onto it. For SIMS analysis, conductive substrates such as gold-coated silicon wafers were used to thaw-mount the tissue on, to prevent charge accumulation during SIMS analysis. The gold coating reduces MS peaks that interfere with lanthanide peaks. Serial sections of the mouse brain were cut and analysed for comparative imaging (Figure 1a and b; Figure 3a and b; Figure 5a and b). All samples discussed in this paper were from two WT mouse brains.

**Figure 1.**
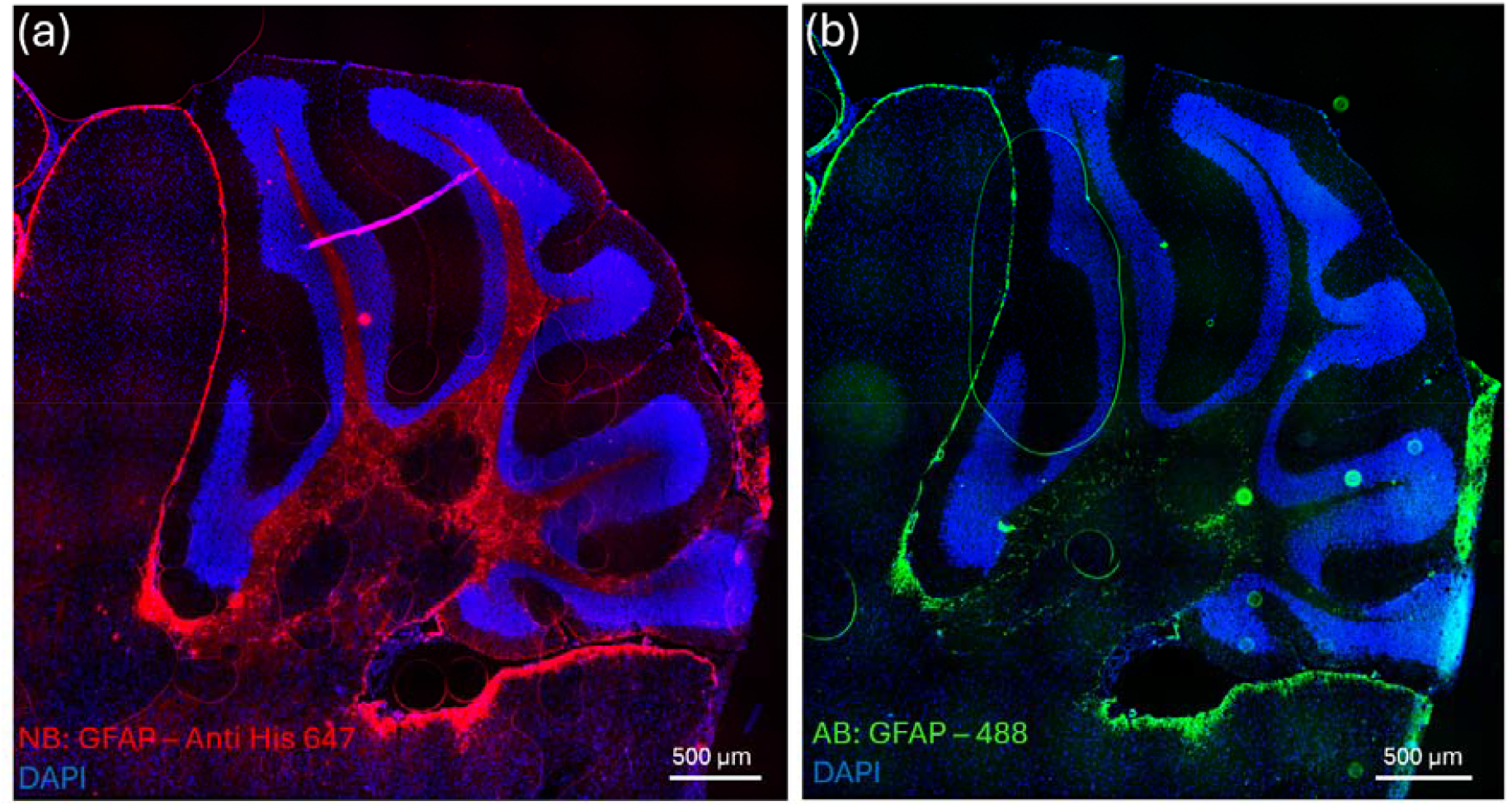
Comparison of NB and AB-488 staining for GFAP in mouse cerebellum using fluorescence imaging. (a) Cerebellar tissue section stained with DAPI (blue) and NB,detected using AB-647 (red). (b) Cerebellar tissue section stained with AB-488 (green). Images were acquired by optical microscopy at 20× magnification.

**Figure 2.**
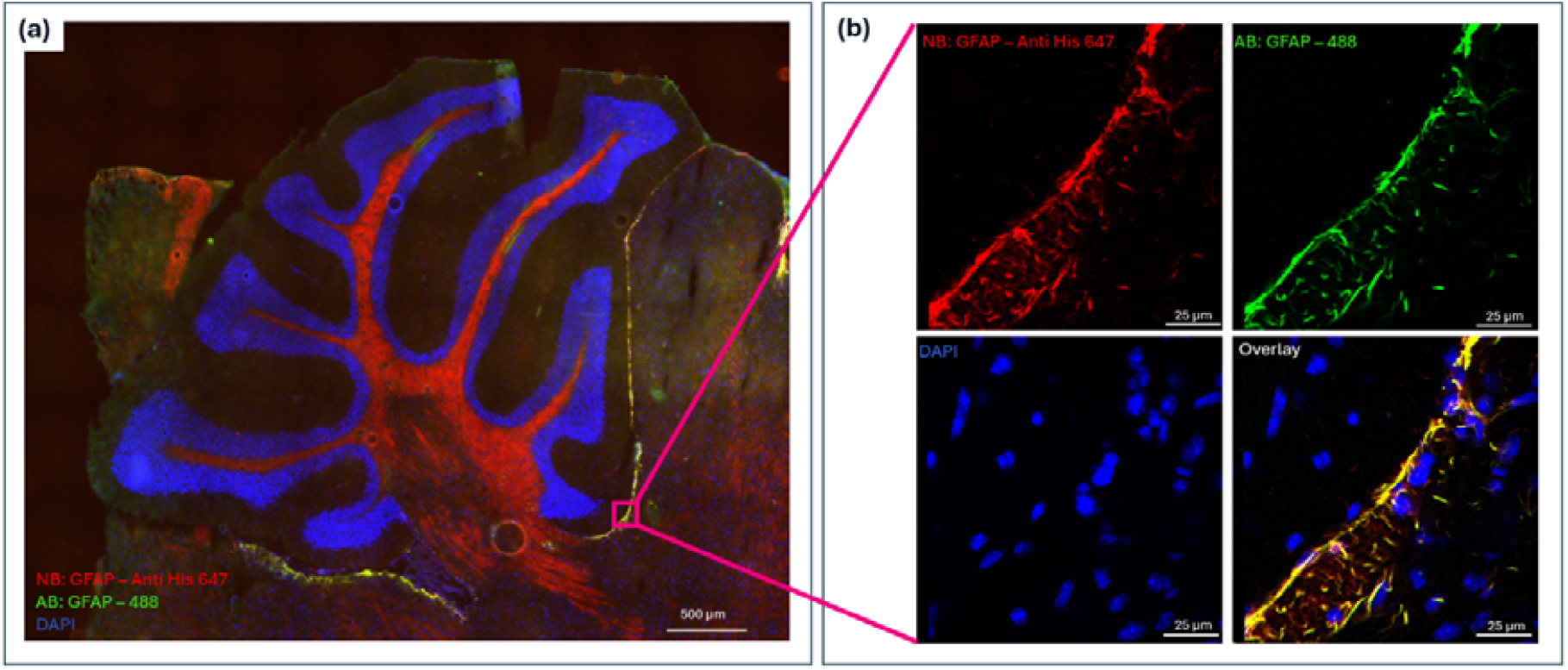
Fluorescence imaging of GFAP staining in mouse cerebellum using both NB and AB-488. (a) Cerebellar tissue stained with DAPI (blue), AB-488 (green), and NB, detected using AB-647 (red). Image acquired by optical microscopy at 20× magnification. (b) Higher-magnification view of the region shown in (a), acquired by confocal microscopy at 40× magnification.

**Figure 3.**
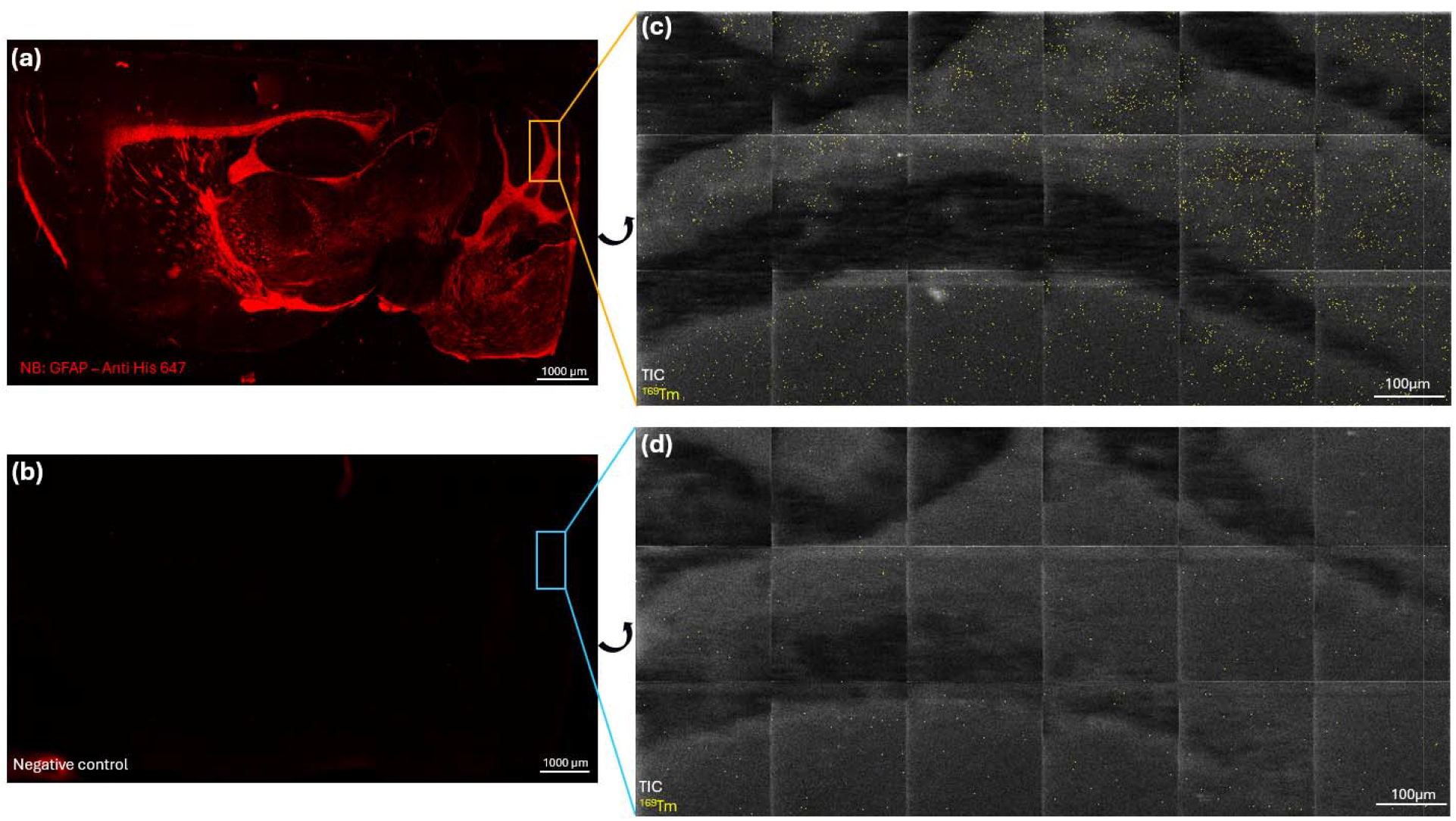
Fluorescence and SIMS imaging of GFAP in mouse cerebellum. (a) Mouse brain tissue section (5 µm thick) stained for GFAP with NB and subsequently AB-647. (b) Negative control without NB or AB-647 staining. (a, b) were acquired by optical microscopy at 20× magnification. (c, d) High resolution SIMS images of the cerebellum, analysed in positive ion mode using a xenon ECR primary ion beam. Panels show the total ion count (TIC; greyscale) and the ^169^Tm signal (yellow) for the NB-stained tissue (c) and negative control (d). The SIMS images (c, d) are rotated by 90 ° relative to the optical microscopy images (a, b).

**Figure 4.**
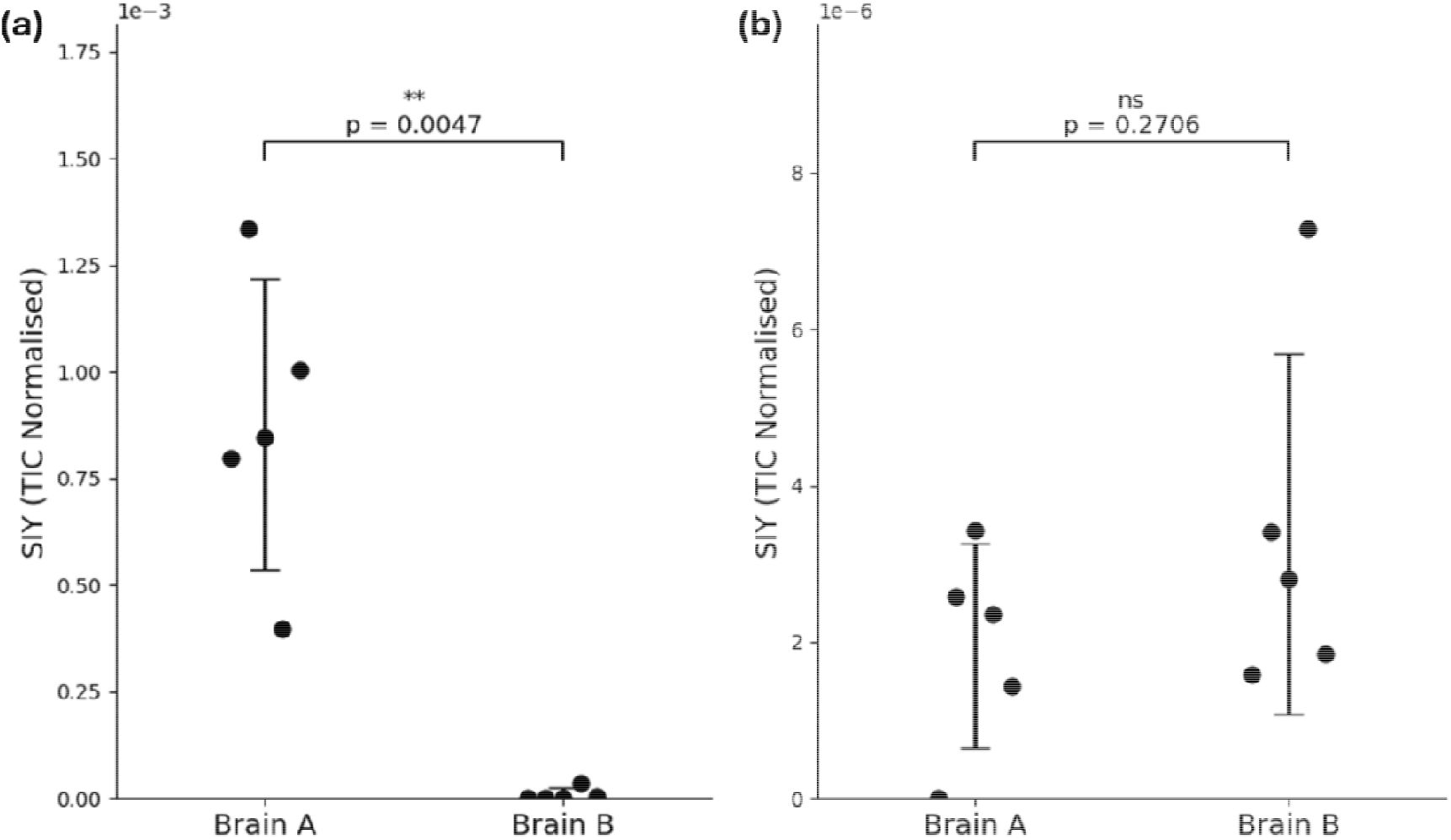
Comparison of the ^169^Tm signal in NB-stained and negative-control mouse brain tissue. (a) Scatter plot showing the ^169^Tm secondary ion yield (SIY), normalised to the TIC, from five randomly selected regions within the cerebellar white matter of the NB-treated mouse brain section (Brain A) and corresponding regions of the negative-control section (Brain B). (b) Equivalent analysis of five randomly selected regions within the cerebellar grey matter of Brain A and corresponding regions of Brain B. Error bars showing the mean ± standard deviation. ** p ≤ 0.01; ns, not significant.

**Figure 5.**
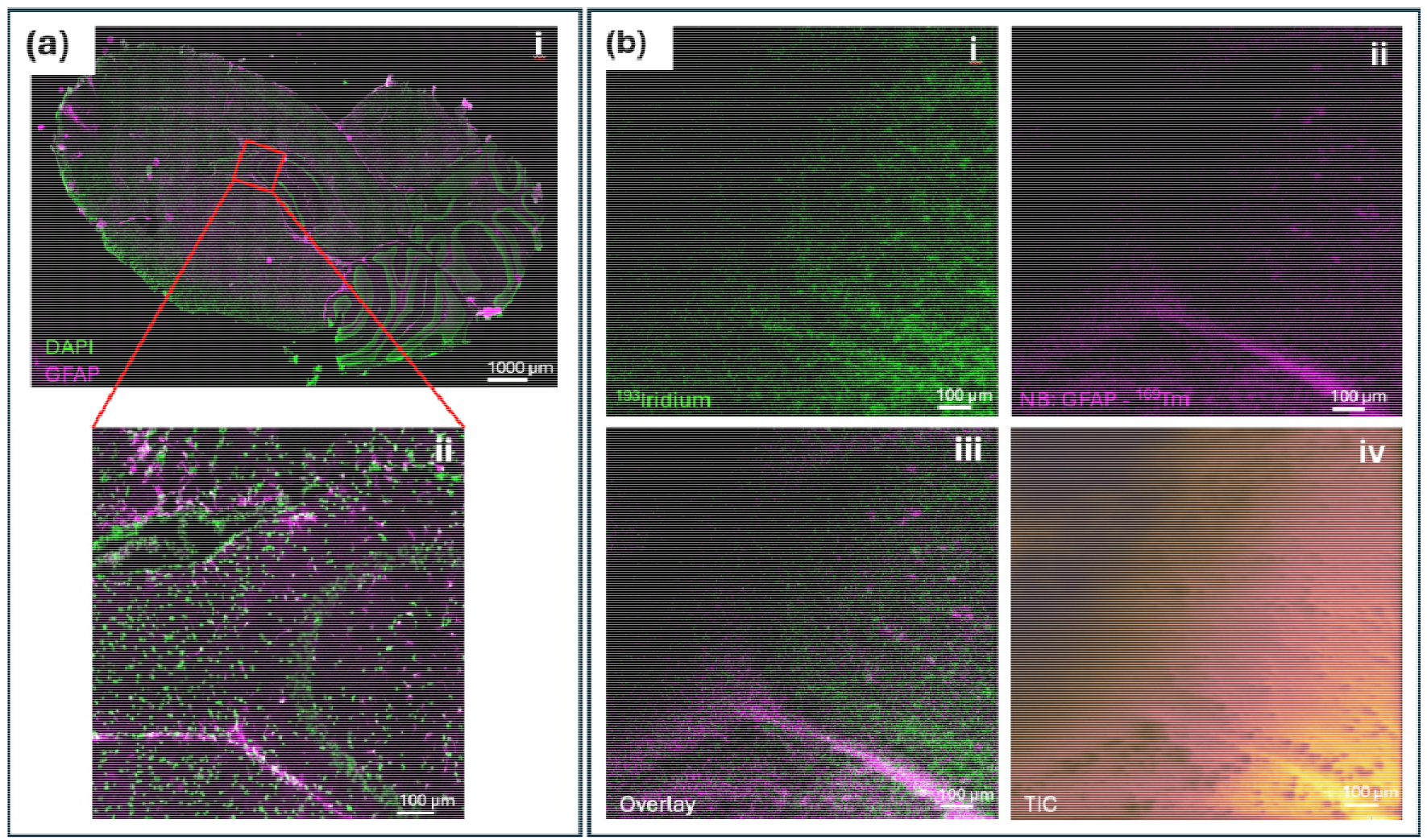
Fluorescence and SIMS imaging of GFAP in mouse hippocampal region. (a) Mouse brain tissue stained for GFAP with AB-488 (purple) and DAPI (green). Image acquired with optical microscopy at 20× magnification. (b) The corresponding region of a serial tissue section analysed by SIMS in positive ion mode using a C_60_^+^ ion primary ion source, stained for GFAP with NB (purple) and for nuclei with ^193^Ir (green).

### Nanobody production and conjugation to lanthanides

A synthetic gene block (IDT Technology) encoding the anti-GFAP nanobody (E9) codon [19], [20] optimised for expression in *E. coli* was inserted into the vector, pPOPINVHHcys [21]. This introduced a C-terminal cysteine for conjugation (Cys-tag) and a hexahistidine tag (His-tag) for detection and purification. The anti-GFAP nanobody was expressed in *E. coli* WK6 cells and purified from the periplasm by a combination of Immobilised Metal Affinity Chromatography (IMAC) and Size Exclusion Chromatography (SEC) as previously described [21].

The commercially available anti-his antibody, Alexa Fluor® 647 Anti-6X His tag® antibody pre-conjugated to the fluorophore 647 (AB-647; ab237337, abcam) was used to tag the anti-GFAP nanobody through the His-tag. The introduced Cys-tag enabled site-specific conjugation of the anti-GFAP nanobody to a Maxpar X8 polymer via the sulfhydryl group of the cysteine residue. The Maxpar X8 polymer was loaded with the lanthanide isotope ^169^thulium (^169^Tm), thereby producing a ^169^thulium-labelled anti-GFAP nanobody (NB). The lanthanide-conjugation to the anti-GFAP nanobody was undertaken using the commercially available kit, Maxpar Antibody Labeling Kit, by Standard Biotools™. The procedure is described in more detail in the paper by Han *et al*. (2018) [22]. After reducing the anti-GFAP nanobody (200 µg) with Tris(2-carboxyethyl)phosphine hydrochloride (TCEP, Sigma-Aldrich), the nanobody was used for conjugation and a final concentration of 0.5 mg/mL of the NB was stored in Phosphate-buffered saline (PBS, Sigma-Aldrich) at 4 °C until further use.

### Immunofluorescent staining

The staining procedure was carried out on samples fixed in -20 °C methanol (Fisher Chemical) and/or 1% paraformaldehyde (Thermo Scientific Chemicals). The commercially available anti-GFAP antibody conjugated to the fluorophore 488 (AB-488; ab302977, abcam) was used for comparing labelling specificity with the NB.

Prior to staining, the tissue sections were washed with phosphate buffered saline (PBS, Sigma-Aldrich Merck) three times for 5 min each. Subsequently, to prevent non-specific binding of the NB, the sections were blocked for 1 h with the blocking buffer consisting of 10% donkey in 0.3% Triton X-100 (Triton X-100, Sigma-Aldrich Merck) in PBS buffer. Next, the sections were incubated overnight at 4 °C with the desired concentration of the NB and AB-488 in blocking buffer (Table S1). The sections were then nuclear stained with cationic nucleic acid intercalator, Cell-ID Intercalator-Iridium isotope ^193^Ir (Fluidigm Corp 201192B), and AB-647, both diluted in PBS, for one hour at room temperature (RT) before washing the sections three times with PBS for 5 min each. Afterwards, the sections were dipped 3 times in deionised water for 15 s each and desiccated at RT for 30 min before stored at 4 °C until SIMS analysis.

Negative control samples went through a similar process to above, only without any staining and were subsequently mounted using a medium containing 4′,6-diamidino-2-phenylindole (DAPI; ab104139, abcam), used for fluorescent staining of the DNA. Samples were coverslip sealed with nail polish for better preservation of the tissue. An outline of the different tissue sections, related staining and control samples is given in Table 1.

**Table 1.**
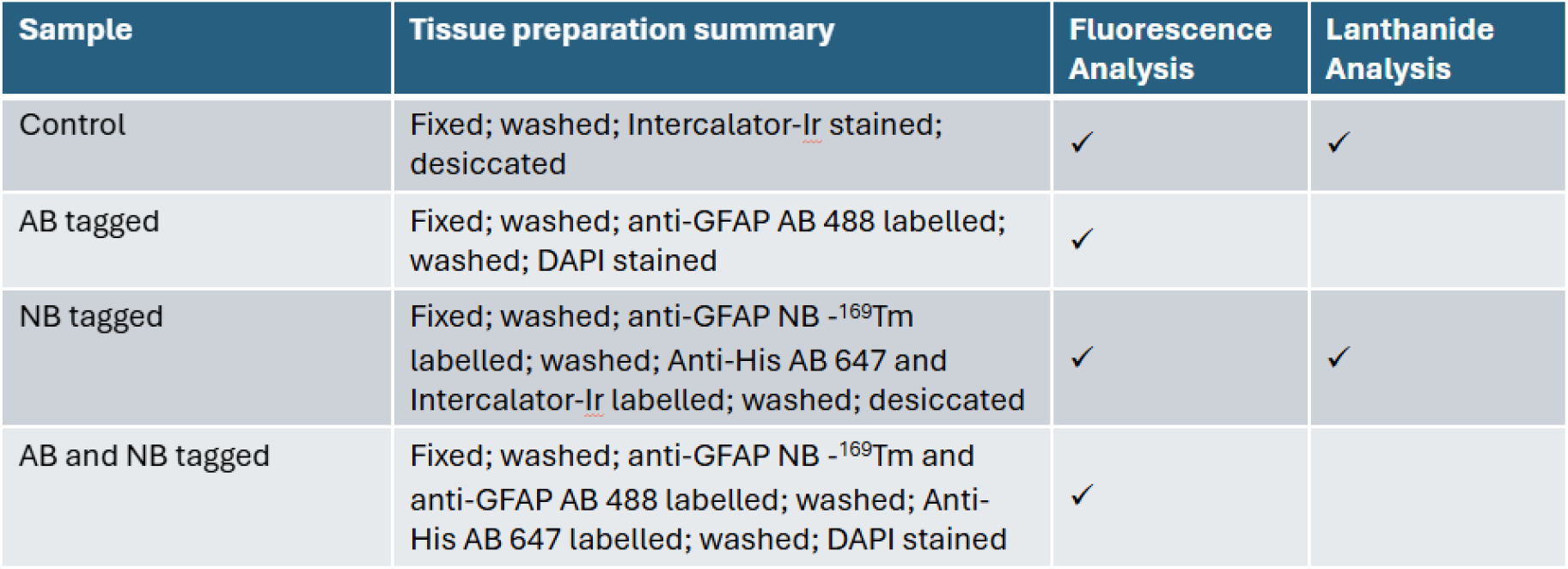
Summary of experimental tissue samples prepared for fluorescent and SIMS analysis.

### Optical microscopy

The stained tissue sections were imaged with the Leica THUNDER Imager Tissue combined with the LMD7 optical microscope using a 5× (0.12 NA), 10× (0.25 NA), and 20× (0.4 NA) objective. Additional imaging was done using the Leica SP8 confocal microscope with a 40× (1.25 NA) oil immersion objective. The wavelengths used to excite the fluorophores were 405 nm for imaging of DAPI, 488 nm for the AB-488, and 647 nm for the AB-647 (which is tagging NBs).

### SIMS analysis

SIMS analysis was performed using a J105 3D Chemical Imager (Ionoptika Ltd., Southampton, UK) [23]. A C_60_^+^ 40 keV ion gun was used to image the sectioned mouse brain tissues stained with the NB in positive ion mode (Figure 5). An ECR ion gun was used to image WT mouse brain sections that have been exposed to NBs (Figure 4c and d). ECR analysis was done in positive mode using an argon or xenon plasma beam. Mass calibration was achieved on ^98^molybdenum and gold, or copper/copper-clusters, depending on the experiment. The thulium lanthanide tag, ^169^Tm, at *m/z* 168.926 (23.15 ppm) was detected along with the Intercalator-Iridium isotope ^193^Ir (*m/z* 192.963, 1.04 ppm) to visualise the localisation of the protein of interest and the nuclei respectively. The region of interest was selected by imaging tissue sections using optical microscopy of the same or adjacent section. The specific parameters used for analysis are summarised in the supplementary information Table S2, Figure S1 and S2. SIMS experimental data was analysed with the Ionoptika Image Analyser Software (version 0.3.3.12) to select single mass channels from the SIMS images or using the software SCiLS™ Lab MVS (version 2026b Pro) by Bruker Corporation.

## Results

### Nanobody production and characterisation

A version of the anti-GFAP nanobody, E9, previously reported by Perruchini et al. [19], [20] incorporating a C-terminal cysteine residue was expressed and purified from *E. coli*. The amino acid sequence of the expressed Cys-tagged anti-GFAP nanobody is: QVQLQVSGGGSVQPGGSLRLSCAASGFTFSNYAMYWLRQAPGKGIEWVSRIGPGGS TEYADSVKGRFTISRDNAKNTLSLQMNSLKPEDTAVYYCAITQQGSGRGQGTQVTV SSGCHHHHHH

Based on the sequence the expected molecular mass of the nanobody is 13.179kDa. Quality assessment of the purified NB by mass spectrometry gave a single peak with a molecular mass of 13.178 kDa confirming the identify and purity of the protein.

### Fluorescent tagging

Fixed serial sections of the cerebellum of WT mouse brains were stained with either the NB (followed by AB-647) or with the AB-488. In both cases the tissue sections were also stained using DAPI. The NB showed corresponding staining patterns to the signal seen with the AB-488, which was most prominent in the white matter (arbor vitae) of the cerebellum, at the outer edge of the inferior colliculus and the medulla (Figure 1). This confirmed that the NB could be used to visualise GFAP with comparable results to the commercially available AB-488.

To further assess the co-localisation of the NB and the conventional AB-488 to GFAP, a WT mouse brain tissue section was simultaneously stained with the NB (with AB-647) and the AB-488, together with DAPI for cell nuclei labelling (Figure 2). Overlap between the NB and AB-488 signals, visualised in yellow, was observed along the outer edge of the medulla and inferior colliculus (Figure 2a). Confocal microscopy at higher magnification of the same tissue section further demonstrated close correspondence between the NB and AB-488 signals within these regions (Figure 2b). These results support the specificity of the NB for GFAP and its ability to reproduce the localisation pattern obtained using a conventional antibody. Interestingly, the fluorescence signal associated with the NB appeared more prominent in the arbor vitae than the AB-488 (Figure 1 and 2). Several factors could contribute to this difference, including biological variation between tissue sections, and differences in the excitation and emission properties, labelling efficiency, and photostability of the fluorophores. Differences in tissue penetration and epitope accessibility may also contribute to the observed staining pattern. Due to their smaller size (∼ 15 kDa), nanobodies diffuse more readily through tissue and access epitopes that may be less accessible to conventional antibodies (∼150 kDa) [18]. However, fluorescence intensity alone cannot distinguish between these factors, and therefore the stronger NB-associated signal should not be interpreted as direct evidence of greater GFAP abundance or enhanced binding properties of the NB. Further studies are required to determine whether nanobodies exhibit higher binding efficiency than conventional antibodies and to assess whether their smaller molecular size provides advantages in tissue penetration and antigen accessibility. Further regions of the brain, such as the hippocampus, also showed co-localisation of the NB and AB-488, specifically in regions surrounding the dentate gyrus and within the polymorphic layer (Figure S3).

### Lanthanide tagging

Adjacent sections of mouse brain tissue were stained with either the NB followed by the AB-647 or without either the NB or AB-647 as a negative control (Figure 3). Other experiments were conducted with only the AB-647 and DAPI as further negative controls (Figure S2). Both samples were imaged by fluorescence microscopy across the whole brain (Figure 3a and b). This showed localisation of the fluorescent tags in the NB and AB-647 stained tissue (Figure 3a) and no observable fluorescent signal in the negative control tissue (Figure 3b). A small region within the cerebellum of these same samples was then imaged with SIMS and the intensity of the ^169^Tm lanthanide peak was mapped (Figure 3c and d). Both Figure 3c and d show a rotated region from Figure 3a and b, respectively. The total secondary ion count shown as greyscale is overlaid with the ^169^Tm peak intensity shown in yellow for the NB labelled GFAP in Figure 3c. This showed an accumulation of the GFAP within the arbor vitae in the NB-stained sample (Figure 3c and 4a) which matched the fluorescence signal and is consistent with results shown in Figure 1 and 2. In contrast, the negative control demonstrated very low ^169^Tm signal for the SIMS image (Figure 3d), and no fluorophore signal in the IF image (Figure 3b). This was further confirmed by semi-quantitative mass spectrometry analysis of peak intensity associated with the presence of ^169^Tm (Figure 4a and b). The relative intensities were sampled in 5 different areas in both the white and grey cerebellar brain matter, in addition to repeating the analysis in both the NB-stained and negative-control brain sections. Statistical analysis was used to assess if ^169^Tm levels were significantly increased after NB staining. By applying the Welch’s independent t-test (Figure 4a and b) a significant difference was observed between the NB-stained and the negative control tissue in the cerebellar white matter (t = 5.68, p = 0.0047), but not the grey matter (t = -1.21, p = 0.2706). ^169^Tm intensities were low in the grey matter in both the NB-stained tissue and the negative control. The signal observed in the NB-treated sample (Figure 3a and c) can be attributed to successful binding of the NB to its target.

In Figure 6 the visualisation of GFAP distribution is compared using conventional antibody-based IF with that obtained by SIMS imaging using lanthanide-conjugated nanobodies. One tissue section was labelled with DAPI as well as AB-488 and imaged by fluorescence microscopy (Figure 5a). A consecutive tissue section was stained with the NB as well as ^193^Ir and imaged by SIMS using a C_60_^+^ ion primary ion source (Figure 5b). The spatial distribution of the AB-488 and DAPI fluorescence signal corresponded closely with those of the ^169^Tm and ^193^Ir signal, respectively, demonstrating a similar localisation of GFAP and cell nuclei between the conventional IF (Figure 5aii) and SIMS (Figure 5biii) imaging methodologies. The close spatial agreement between the conventional IF and SIMS images supports the successful localisation of GFAP using the lanthanide-conjugated nanobody. These results demonstrate the feasibility of lanthanide-conjugated nanobodies as a novel labelling approach for protein localisation by SIMS.

Importantly, these findings establish the feasibility of lanthanide-conjugated nanobodies as an alternative to conventional antibody-based labelling strategies for SIMS imaging. This represents a significant step towards expanding the range of molecular probes available for mass spectrometry imaging, highlighting the potential of nanobodies for high-resolution protein localisation and future multiplexed tissue imaging applications.

## Conclusion

This study demonstrated the potential of lanthanide-conjugated nanobodies as probes for protein localisation using SIMS imaging. The anti-GFAP nanobody showed comparable localisation to a commercially available anti-GFAP antibody in fluorescence microscopy, with clear co-localisation observed in several regions of the mouse brain. Furthermore, conjugation of the nanobody to a ^169^Tm-containing polymer enabled detection of the target-associated lanthanide signal by SIMS, with the observed distribution consistent with fluorescence imaging and substantially reduced background signal in unstained control tissue. These findings provide evidence that nanobodies can be successfully adapted for lanthanide-based mass spectrometry imaging. Their smaller size and favourable stability may offer advantages over conventional antibodies, potentially improving tissue penetration and spatial localisation. However, one of the main limitations of nanobody-based imaging is the limited commercial availability of target-specific nanobodies compared with conventional antibodies. This may restrict their broader application and necessitate the development and production of custom nanobodies for specific antigens. Additionally, further optimisation and quantitative comparison with antibody-based tags are required. In conclusion, this work establishes a promising approach for expanding lanthanide multiplexing to nanobody-based tags. Such tags could ultimately enhance the spatially resolved analysis of proteins and support multimodal and multi-omics imaging approaches.

## Acknowledgements

The authors thank the Rosalind Franklin Institute (RFI) for the resources and funding that supported this work. We also thank members of the RFI Technology Innovation Challenges, “Integrated Chemical Imaging in Cells and Tissues”. In particular, we thank Dorcas Cheung for assistance with the immunofluorescence methodology. We are grateful to Dr Logan Mackay for the help and knowledge around instrumental issues. We also acknowledge Ionoptika Ltd. for their technical support and assistance. The Rosalind Franklin Institute is funded by UK Research and Innovation through the Engineering and Physical Sciences Research Council.

## Supplementary information

**Table S1.** Dilution of NB, AB-488, AB-647, and 193Ir in blocking buffer or PBS as indicated in methodology section. All stock concentrations were 0.5 mg/mL.

| Dilution | Figure |
| --- | --- |
| NB (1:200) | 1, 2 |
| NB (1:50) | 3 |
| NB (1:250) | 5 |
| AB-488 (1:200) | 1, 2 |
| AB-488 (1:400) | 5 |
| AB-647 (1:500) | 1, 2, 3 |
| <sup>193</sup>Ir (1:1000) | 5 |

**Table S2.** Experimental parameters for all analyses done with the SIIMS. The spot size (spatial resolution) was measured using the 20/80 percent method.

| Sample | Ion source | Total ion dose (ions/cm<sup>2</sup>) | Number of pixels | Area analysed (µm × µm) | Spatial resolution (µm) |
| --- | --- | --- | --- | --- | --- |
| Figure 4<br>Positive control | ECR - Xe<sup>+</sup> | 2.21 × 10<sup>15</sup> | 128 | 1152 × 576 | 1.99 |
| Figure 4<br>Negative control | ECR - Xe<sup>+</sup> | 2.21 × 10<sup>15</sup> | 128 | 1152 × 576 | 1.99 |
| Figure 5 | C<sub>60</sub><sup>+</sup> | 3.82 × 10<sup>13</sup> | 512 | 1200 × 1200 | 2.28 |

**Figure S1.**
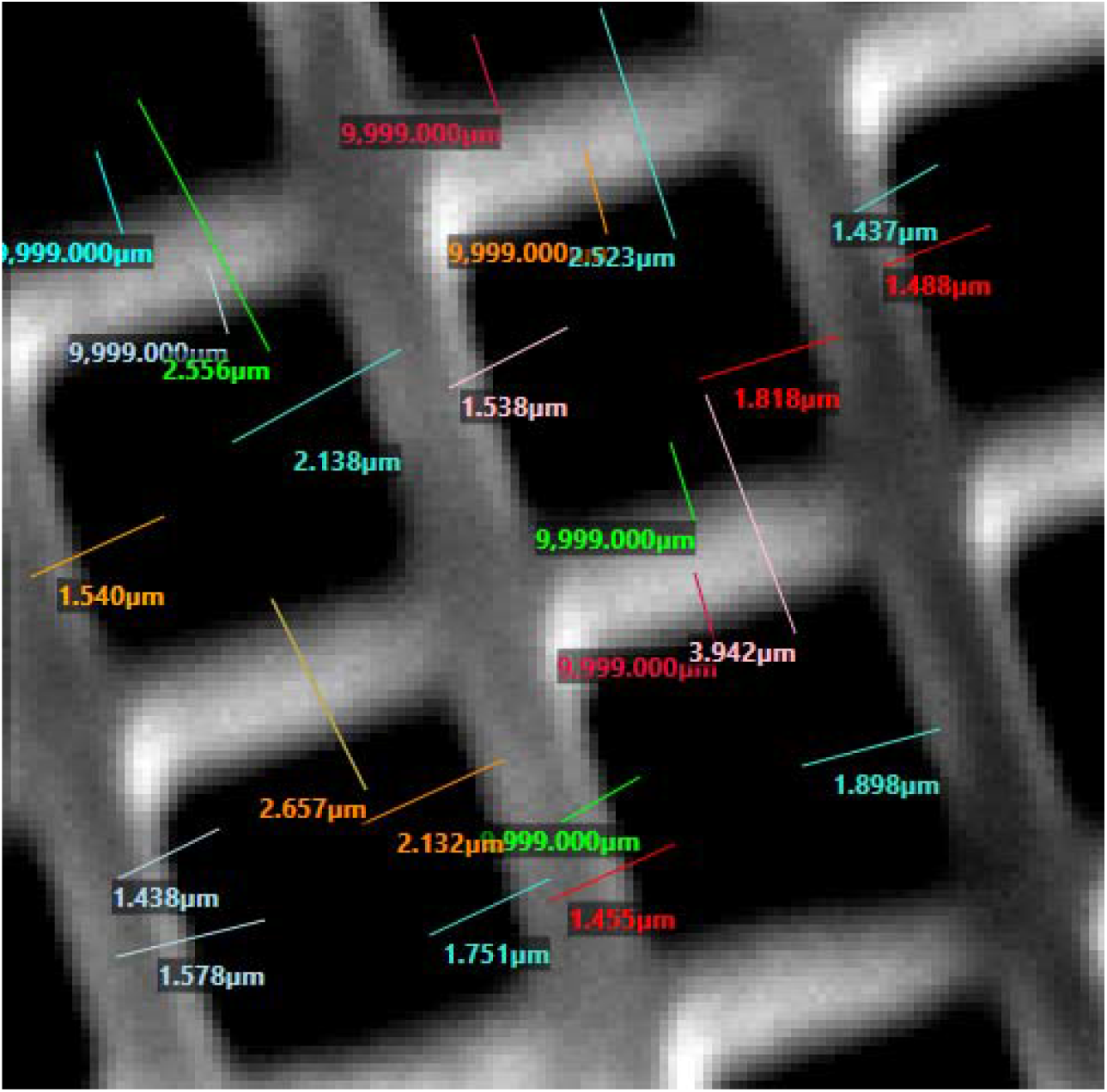
ECR beam spot size used for imaging of Figure 4c and d. Mean of spot size measurements ± standard deviation (excluding 9,999.00 µm measurements due to software errors): 1.99 µm ± 0.67.

**Figure S2.**
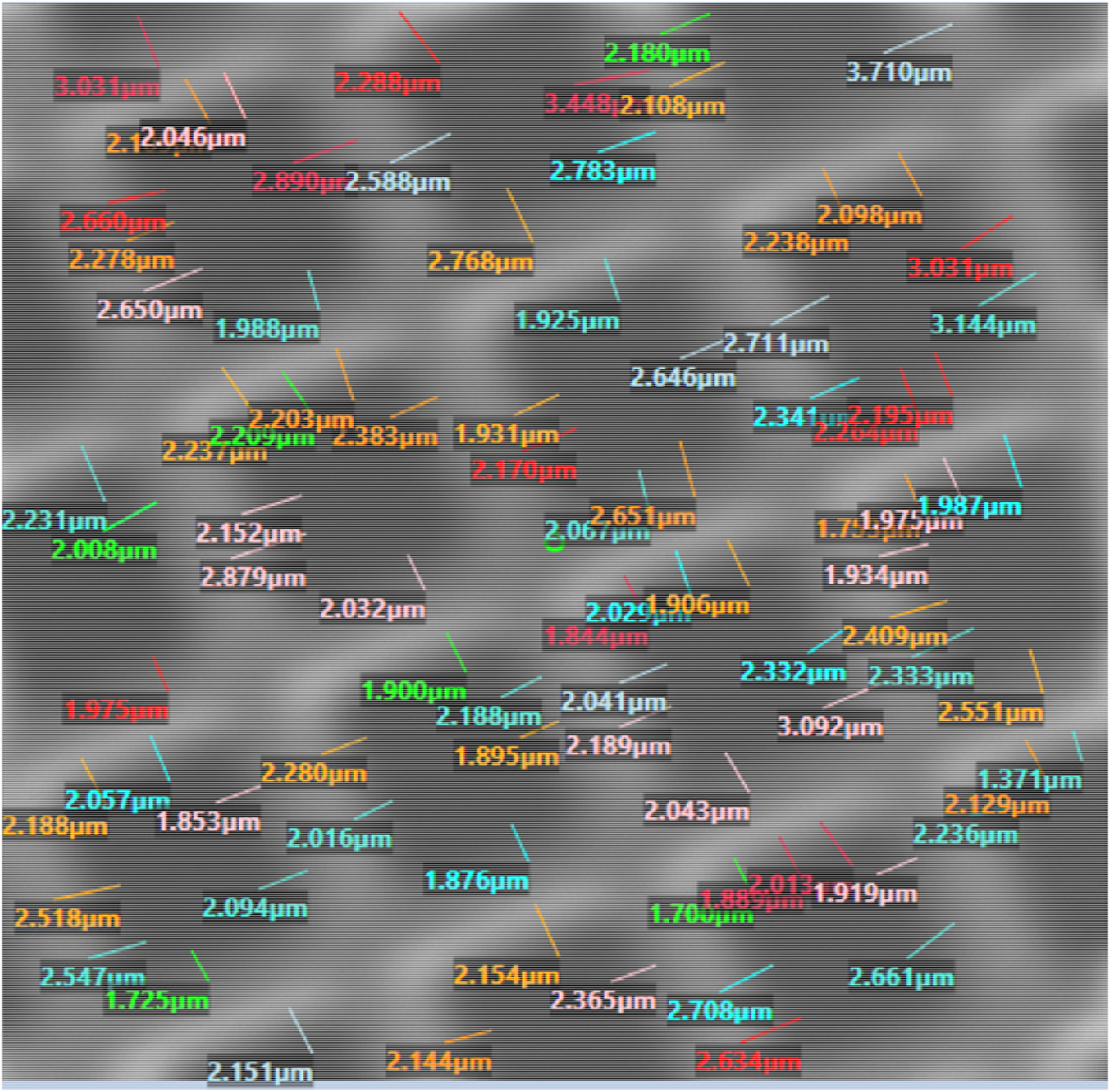
C_60_ beam spot size used for imaging of Figure 6b. Mean of spot size measurements ± standard deviation: 2.28 µm ± 0.41.

**Figure S3.**
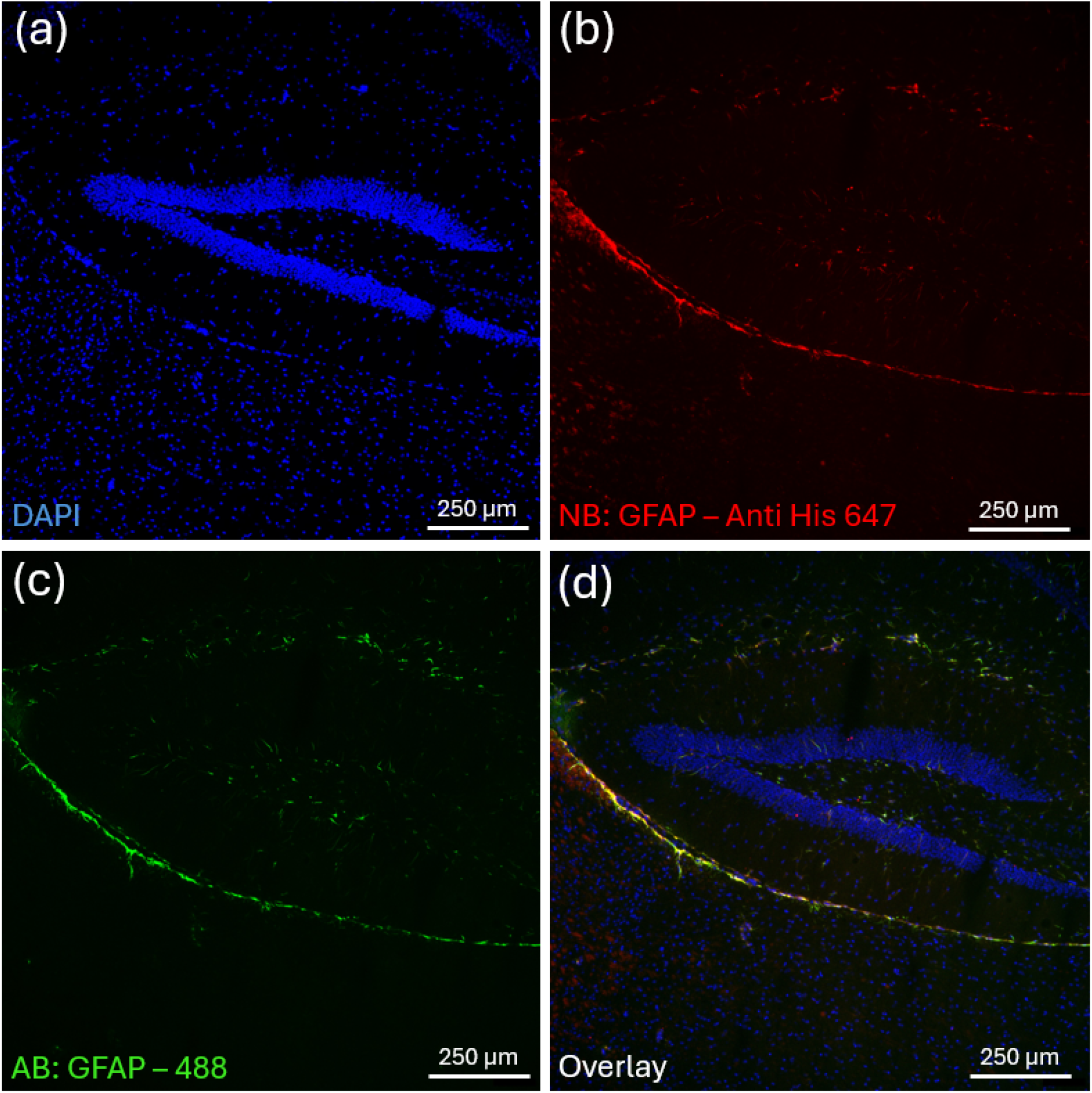
Fluorescence imaging of GFAP staining in mouse hippocampus using both NB and AB-488. Hippocampal tissue stained with (a)DAPI (blue), (b) NB detected using AB-647 (red), (c) AB-488 (green), and (d) showing the overlay. Image acquired by using the Leica THUNDER Imager Tissue combined with the LMD7 optical microscope at 20× magnification.

**Figure S4.**
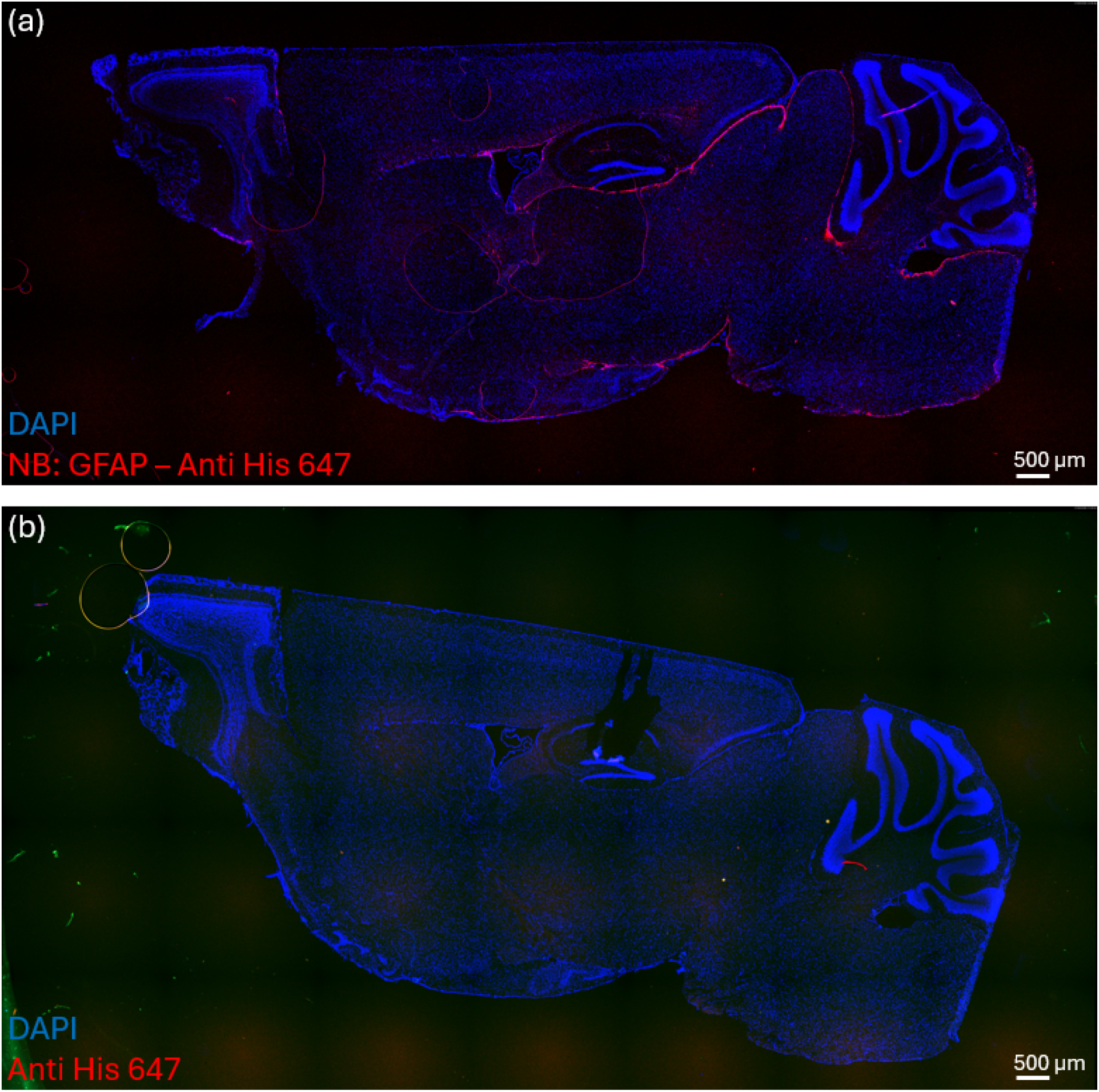
Fluorescence imaging of wt mouse brain. (a) stained for DAPI (blue) and anti-GFAP nanobody with a His-tagged antibody pre-conjugated to fluorophore 647 (red), (b) negative control with only the His-tagged antibody pre-conjugated to fluorophore 647. (a) and (b) are imaged with a 5x magnification using the Leica THUNDER Imager Tissue combined with the LMD7 optical microscope.

